# Transfer Learning for Predicting Virus-Host Protein Interactions for Novel Virus Sequences

**DOI:** 10.1101/2020.12.14.422772

**Authors:** Jack Lanchantin, Tom Weingarten, Arshdeep Sekhon, Clint Miller, Yanjun Qi

## Abstract

Viruses such as SARS-CoV-2 infect the human body by forming interactions between virus proteins and human proteins. However, experimental methods to find protein interactions are inadequate: large scale experiments are noisy, and small scale experiments are slow and expensive. Inspired by the recent successes of deep neural networks, we hypothesize that deep learning methods are well-positioned to aid and augment biological experiments, hoping to help identify more accurate virus-host protein interaction maps. Moreover, computational methods can quickly adapt to predict how virus mutations change protein interactions with the host proteins.

We propose DeepVHPPI, a novel deep learning framework combining a self-attention-based transformer architecture and a transfer learning training strategy to predict interactions between human proteins and virus proteins that have novel sequence patterns. We show that our approach outperforms the state-of-the-art methods significantly in predicting Virus–Human protein interactions for SARS-CoV-2, H1N1, and Ebola. In addition, we demonstrate how our framework can be used to predict and interpret the interactions of mutated SARS-CoV-2 Spike protein sequences.

**Availability:** We make all of our data and code available on GitHub https://github.com/QData/DeepVHPPI.

**ACM Reference Format:** Jack Lanchantin, Tom Weingarten, Arshdeep Sekhon, Clint Miller, and Yanjun Qi. 2021. Transfer Learning for Predicting Virus-Host Protein Interactions for Novel Virus Sequences. In *Proceedings of ACM Conference (ACM-BCB)*. ACM, New York, NY, USA, 10 pages. https://doi.org/??

## 1 INTRODUCTION

A protein-protein interaction (PPI) denotes a critical process where two proteins come in contact with each other to carry out specific biological functions. Virus proteins, such as those from the 2019 novel coronavirus, also known as SARS-CoV-2, interact with human proteins to infect the human body, and ultimately overtake physiological functions (e.g., alveolar gas exchange). Accordingly, protein-protein interactions are often the subject of intense research by virologists and pharmaceutical scientists. Knowing and understanding which host proteins a virus with a novel sequence pattern may interact with is crucial. Such discoveries will expedite our understanding of virus mechanisms and may aid in the development of vaccines, diagnostics, therapeutics, and antibodies.

We aim to infer possible interactions between all host proteins and a novel virus protein or a novel variant. This setup is beneficial for three reasons. First, our model can predict an initial set of interactions if experiments have not yet been done. Second, our model can expand the initial set of experimental interactions, resulting in a more complete interactome. Finally, such computational models enable us to test hypotheses such as the effect of mutations.

While protein-protein interaction information is expensive to obtain, protein sequence information is cheap and fast. In this paper, we propose a deep neural network (DNN) based framework, DeepVHPPI, to predict protein interactions between virus proteins and host proteins using sequence information alone. DeepVHPPI includes two key designs: (1) Motivated by the evidence that co-occurring short polypeptide sequences between interacting protein partners appear to be conserved across different organisms [47], we introduce a novel DNN architecture to learn short sequence patterns, or “protein motifs” via self-attention based deep representation learning. (2) since virus-host PPI data is limited, we propose a transfer learning approach to pretrain the network on general protein syntax and structure prediction tasks. The objective of this transfer learning approach is to improve generalization on the task of predicting protein-protein interactions involving novel virus proteins with unseen sequences.

In summary, we make the following contributions:

- We introduce the DeepVHPPI, a novel deep neural framework for protein sequence based Virus–Host PPI prediction for novel virus proteins or virus proteins with novel variants.
- DeepVHPPI combines a self-attention based transformer architecture and transfer learning for PPI prediction in the context of novel virus sequences (where no previous interactions are known).
- We evaluate DeepVHPPI with validated interactions on Virus–Host PPIs across three virus types: SARS-CoV-2, H1N1 and Ebola datasets. We show that DeepVHPPI outperforms the previous state-of-the-art methods, as well as provide an analysis of SARS-CoV-2 Spike protein mutations.

## 2 BACKGROUND AND TASK FORMULATION

Proteins are biomolecules that are comprised of a linear chain of amino acids. This allows them to be described by a sequence of tokens (each token is one amino acid (AA)). The dictionary of possible tokens contains 20 standard AAs, two non-standard AAs: selenocysteine and pyrrolysine, two ambiguous AAs, and a special character for unknown (i.e. missing) AAs. In other words, we can represent proteins as strings built from a dictionary *V* of size | *V*| = 25. We represent a protein x as a sequence of characters *x*_1_, *x*_2_, …, *x*_*L*_. Each character *x*_*i*_ is one possible amino acid from *V*. Proteins rarely act in isolation but instead interact with other proteins to perform many biological processes. This is referred to as a protein-protein interaction (PPI).

### Task Formulation

Viruses infect a host through Virus-Host PPIs. Therefore, predicting which host proteins a virus protein will bind to is a key step in understanding viral pathogenesis^1^ and designing viral therapies. Identifying virus-host PPI interactions can be formulated as a binary classification problem: “given virus protein sequence **x**^*q*^ and host protein sequence **x**^*k*^, does the pair interact or not?”. Fig. 1 gives a visual representation of the types of PPIs that we consider with our model. that occur within the human body. There are three types of proteins in this diagram: Host (human) proteins, previously known virus proteins, and a novel virus protein. As shown with solid lines, there are a set of known interactions between a pairs of host proteins as well as between known virus and host proteins. We consider known host-host interactions, known virus-host interactions, and unknown virus-host interactions. Fig. 1 visualizes the case of predicting unknown interactions between human proteins and SARS-CoV-2 proteins, given what is known about interactions with proteins from HIV and Zika. Our target task is to predict all possible unknown sets of interactions between the novel virus and host proteins, as shown with a dashed line. This formulation motivates the use of transfer learning because we want to transfer the learned interactions from known viruses to a novel virus. Specifically, we’re concerned here with proteins from novel viruses, which is different than a novel protein from a known virus in our training set

**Figure 1:**
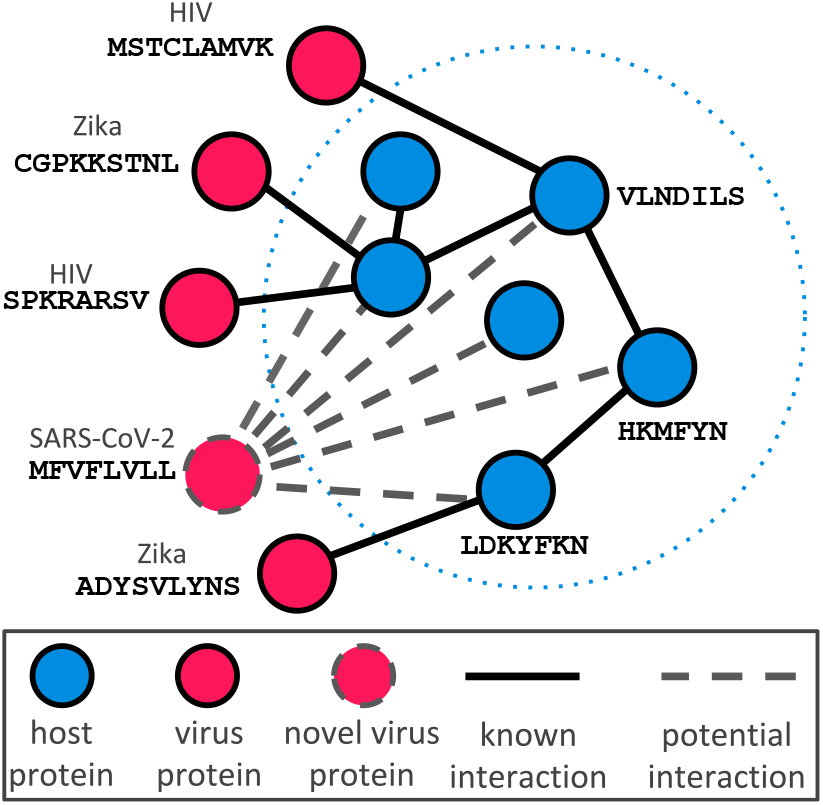
Virus-Host Protein-Protein Interactions (PPI). Overview of our task, where there is a set of previously known protein-protein interactions. Our goal is to predict all possible Virus–Human interactions for a novel virus protein, such as SARS-CoV-2.

### Biological Experiments to Detect PPIs

It remains difficult to accurately uncover the full set of protein-protein interactions from biological experiments. Traditionally, PPIs have been studied individually through the use of genetic, biochemical, and biophysical techniques such as measuring natural affinity of binding partners in-vivo or in-vitro [46]. While accurate, these small-scale experiments are not suitable for full proteome analyses [70]. This is because, for example, there are roughly | *P*_*h*_| ≈ 20, 000 different human proteins, and | *P*_*ν*_| ≈ 26 different virus proteins (in SARS-CoV-2, not considering mutated variants), so the potential search space of V–H interactions is | *Pν*| × | *P*_*h*_| = 0.5M. This number can grow significantly larger when you consider virus protein variants.

High-throughput technologies, such as yeast two-hybrid screens (Y2H) [19] and Affinity-purification–mass spectrometry (AP-MS) [21, 27], are chiefly responsible for the relatively large amount of PPI evidence. Notably, the first experimental study for SARS-CoV-2 interactions used AP-MS [21]. However, these datasets are often incomplete, noisy, and hard to reproduce [64]. The resulting low sensitivity of high-throughput experiments is unfavorable when trying to fully understand how the virus interacts with humans.

### Past Machine Learning based PPI Prediction Studies

Most previous computational methods to predict PPIs have focused on within-species interactions [7, 20, 22, 23, 36, 47, 61, 67, 69]. These methods do not easily generalize to cross-species interactions (e.g., V–H) [68]. Few methods have attempted to predict cross-species protein interactions between humans and a novel virus [68, 71]. Furthermore, previous methods operating at the sequence level do not use structural information from previously known proteins to aid learning [18]. By training virus–host interactions from a variety of viruses and leveraging prior structural information, our proposed model, DeepVHPPI, allows us to predict the host interactions of an unseen virus protein.

When designing machine learning models to predict V–H PPIs, two challenges stand out: (1) Existing sequence analysis tools focus on global alignment patterns while PPIs mostly depend on local binding motif patterns. (2) It is especially difficult for virus proteins that are unknown or are new variants, since there is limited or no experimental interaction data for those sequences. This requires the machine learning model to transfer knowledge from one domain (previously known sequences) to a new domain (novel virus sequences). We argue that this is a realistic task when an unknown virus is newly discovered. In other words, we want to rapidly predict all the host interactions of a newly sequenced virus protein. In this work, we propose a deep learning based pipeline to combine neural representation learning and transfer learning for solving the listed obstacles. Recent literature shows some successful transferability of large scale deep learning models on protein sequences to multiple downstream tasks [52, 57]. To the authors’ best knowledge, we are the first to adapt the self-attention based transfer learning to the virus-host PPI prediction task.

## 3 PROPOSED DNN FRAMEWORK FOR VIRUS HOST PPI PREDICTION: DEEPVHPPI

We denote each amino acid in a protein sequence as x_CLS_, x_1_, x_2_, …, x_*n*_, where x_*i*_ ∈ ℝ^| *V* |^ denotes a one-hot vector and x_CLS_ is a special classification token. Given virus protein x^*a*^ ∈ℝ^*n*×| *V* |^ and human protein x^*b*^ ∈ℝ^*n*×| *V* |^, the goal of DeepVHPPI is to predict the interaction likelihood *ŷ* of the pair of proteins. In this section, we explain the DeepVHPPI architecture, as shown in Fig. 2, which maps a protein sequence x ∈ℝ^*n*×| *V* |^ to representation z ∈ℝ^*n*×*d*^. In section 3.1, we introduce the Transformer module which maps a protein sequence x to a hidden representation, z. In section 3.2, we introduce the classification module which takes as input both the virus protein hidden representation, z^*a*^, and the host protein hidden representation, z^*b*^, and outputs the likelihood that the two proteins interact.

**Figure 2:**
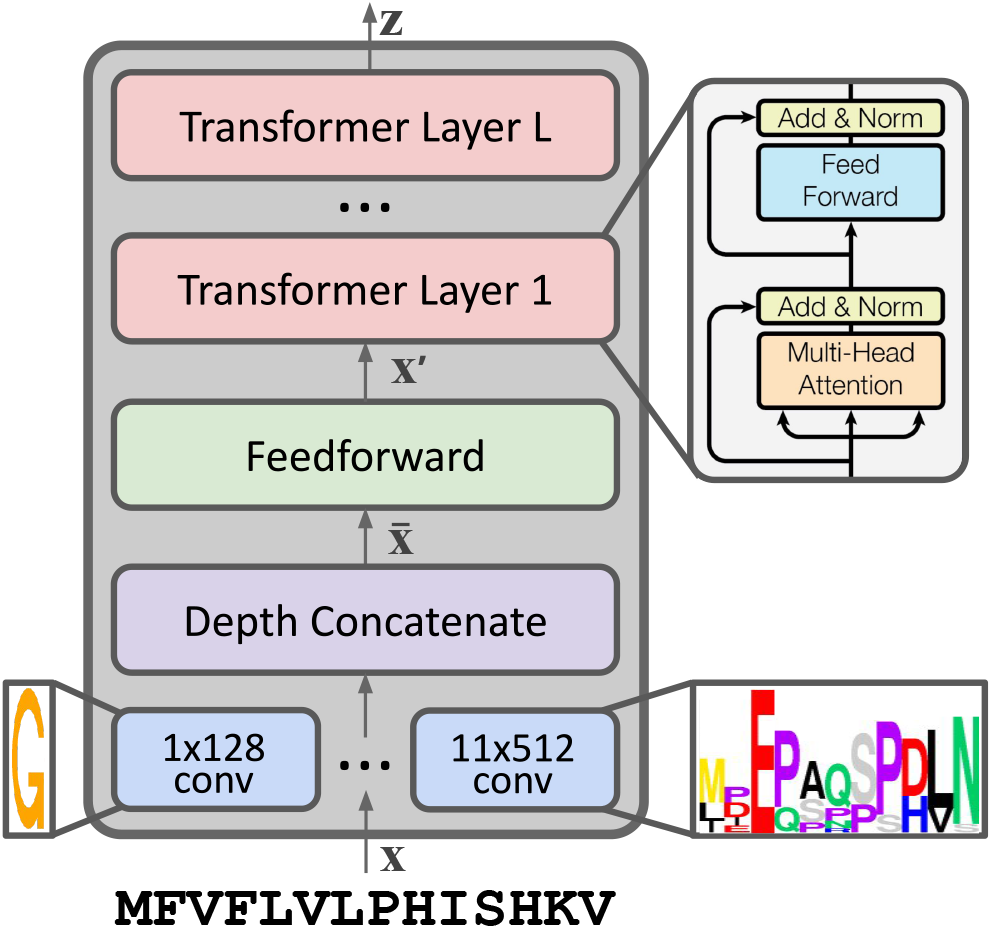
DeepVHPPI Architecture. A one-hot encoded sequence x gets input to the convolutional layers to find protein “motifs”. The convolution outputs are then concatenated along the depth dimension and input to a feedforward layer. Finally, several Transformer encoder layers [63] model the dependencies between the learned convolutional motifs, producing a final representation z. The representation can then be used for any arbitrary classifier layer to predict protein properties.

### 3.1 Transformer Layers to Learn Representations of Protein Sequences

Transformers [63] have obtained state-of-the-art results in many domains such as natural language [17], images [51], and protein sequences [57]. A Transformer encoder layer is a parameterized function mapping input token sequence ∈x ℝ^*n*×*d*^ to z ∈ℝ^*n*×*d*^. At a high level, a Transformer encoder layer “transforms” the representations of input tokens (e.g., amino acids) by modeling dependencies between them in the form of attention. The importance, or weight, of token ***x*** _*j*_ with respect to ***x***_*i*_ is learned through attention. Each Transformer encoder layer performs the following computation on input x:

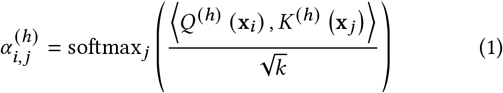

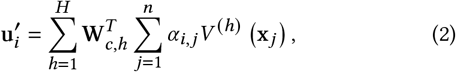

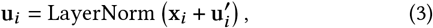

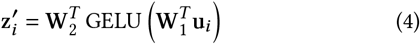

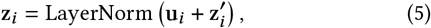

where 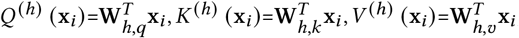 and **W**_*h,q*_, **W**_*h,k*_, **W**_*h,ν*_ ∈ ℝ^*d*×*k*^, **W**_1_ ∈ ℝ^*d*×*m*^, **W**_2_ ∈ ℝ^*m*×*d*^, **W**_*c,h*_ ∈ ℝ^*k*×*d*^. *H, k, m*, and *d* are hyperparameters where *H* is the total number of Transformer “heads”, and *k, m*, and *d* are weight dimensions. GELU is a nonlinear layer [26], and LayerNorm is Layer Normalization [4]. The final z representation after *L* layers is the output of the Transformer encoder, which can then be used by a classification layer.

### Convolutional Layers to Extract Local Motif Patterns

Protein sequences have short, local patterns known as sequence motifs, that have been a major bioinformatics tool for years [54]. If we view amino acids as the protein analog of natural language characters, motifs are analogous to words. In particular, virus proteins that successfully mimic host proteins and interact with other host proteins often display similar motifs to the target of mimicry [16]. To take advantage of these patterns, we introduce an architecture variant that stacks convolutional layers and transformer layers. The key contribution of this variation is to automatically learn sequence motifs via convolutional layers (motif module), and compose local patterns together via deeper transformer layers. Our motif module utilizes different length convolutional filters to find motifs directly from sequence end-to-end.

Specifically, we apply six temporal convolutional filters of sizes {(1×128), (3×256), (5×384), (7×512), (9×512), (11×512)} to the onehot encoded protein sequence ***x*** ∈ ℝ^∈*L*×| *V* |^, where the first number of each filter is the width and the second number is the depth. Each filter is zero-padded to preserve the original sequence length. We depth-concatenate the output of the convolutional, producing 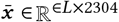. 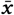 is fed to a Feedforward layer to produce a *L* × *d* matrix 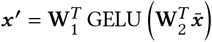 Finally, to encode positional information we add sinusoidal position tokens [63] to the ***x*** ′ matrix. This output ***x*** ′ is used as input to a Transformer encoder.

Using several convolutional filters of varying size allows the model to learn a diverse set of motifs. Specifically, in our implementation, the set of filters allows the model to learn 2304 unique motifs of varying lengths. DeepVHPPI is illustrated in Fig. 2 (left).

### 3.2 Classification Layer to Predict V–H Protein Interactions

The Convolutional and Transformer layers map virus protein sequence x^*a*^ to z^*a*^ and host sequence x^*b*^ to z^*b*^. The final layer of DeepVHPPI is to predict the interaction likelihood of x^*a*^ and x^*b*^. We first obtain a single vector representation of each protein using the the classification token outputs from the Transformer, 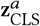 and 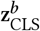. In other words, each protein is fed into the shared Transformer model, and outputs an independent vector. Using these representations, we can predict *ŷ*, the likelihood that the two proteins interact with one another:

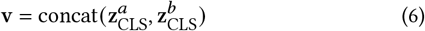

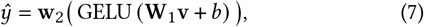

where **W**_1_ ∈ ℝ^2*d*×*d*^, and w_2_ ∈ ℝ^1×*d*^ are projection matrices, and v ∈ ℝ^2*d*×1^ is a concatenation of the two protein representations. GELU is a nonlinear activation layer [26]. The virus sequence is always the first *d* weights in the concatenated representation, so the classifier is not invariant to the protein ordering. *ŷ* is then fed through a sigmoid function to obtain the interaction probability.

### 3.3 Proposed Training: Transfer Learning for Virus–Human PPI Prediction

Our proposed DeepVHPPI network allows us to predict the interaction likelihood of two proteins given only sequence information of each protein. With this framework, we are faced with several difficulties in order to predict Virus-Host interactions. First, there is limited Virus-Host PPI data available to train on. In particular, there are few or no interactions known for novel virus protein sequences. Second, protein structure information is important for accurate PPI prediction [13] Using sequence features alone may not be sufficient for predicting certain interactions, but obtaining structure for novel proteins is a slow process. Both of these obstacles require a model that can generalize from knowledge learned in related protein prediction tasks. To this end, we introduce a “transfer learning” three-step training procedure. This involves pretraining the Convolutional and Transformer layers (Section 3.1) of DeepVHPPI, before fine-tuning the entire network (Section 3.1 and 3.2) on the PPI task.

The first step is to pretrain the DeepVHPPI using Masked Language Model (MLM) pretraining in order to learn generic representations from unlabeled protein sequences; the second step is to further pretrain the network using Structure Prediction (SP) to learn 3D structural representations; and the third step is to finetune the network on the Virus–Host PPI data for previously known viruses. Pretraining the network allows it to learn representations that transfer well to the PPI task for novel (i.e., unseen) virus sequence. An overview of our proposed training procedure is shown in Figure 3, and we explain each training step below. Each task uses a task-specific classifier, shown as MLM, SS, CT, RH, and PPI in Figure 3. In other words, the DeepVHPPI is shared between all tasks, but the classifier layers are not.

**Figure 3:**
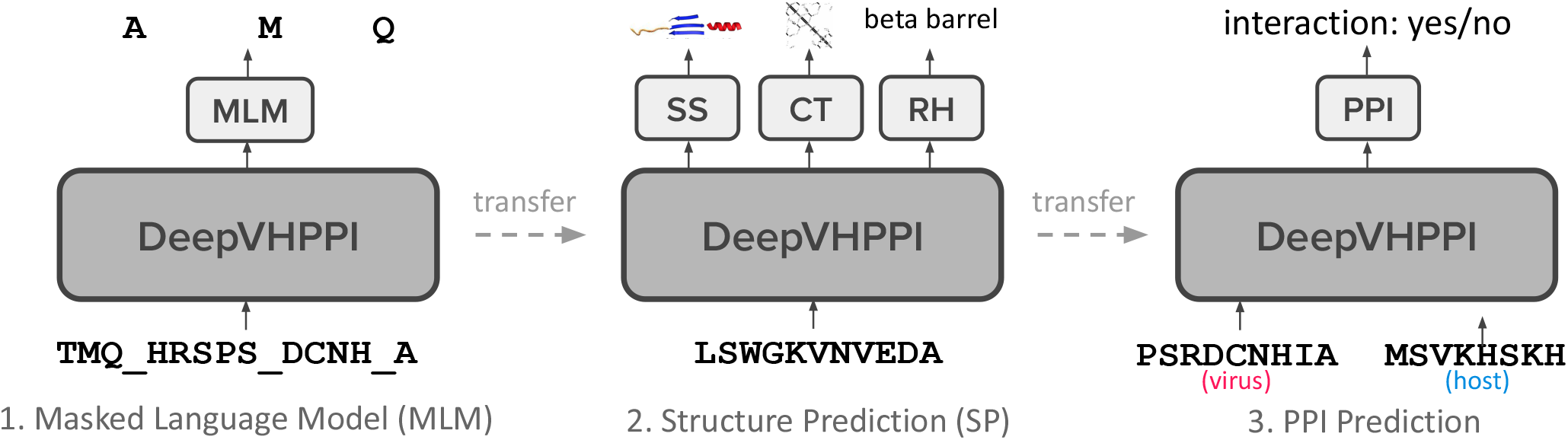
Transfer Learning Framework for DeepVHPPI. First, we pretrain the network on the Masked Language Model (MLM) task from a large repository of unlabeled protein sequences. Second, we further pretrain the network on a set of Structure Prediction (SP) tasks including secondary structure (SS), contact (CT), and remote homology (RH). Finally, we fine-tune the network on the protein-protein interaction (PPI) prediction task. The base DeepVHPPI shown as the large dark grey block is shared across all tasks, and each task uses its own classifier, shown as small light grey blocks.

#### 3.3.1 MLM: Masked Language Model Pretraining

Recent literature on learning self-supervised representations of natural language have shown that pretraining using self-supervised and supervised methods encourages the model to learn semantics about the input domain that can help prediction accuracy on new tasks [8, 12, 17, 37, 44]. In order to learn basic protein semantics and syntax, leveraging large databases of protein sequences is paramount. Masked language model (MLM) training is a self-supervised technique to allow a model to build rich representations of sequences. Specifically, given a sequence x, the MLM objective optimizes the following loss function:

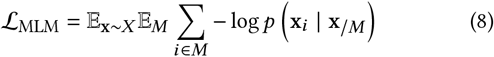

where M is a set of indices to mask, replacing the true character with a dummy mask character. The total loss is a sum of each masked character’s negative log likelihood of the true amino acid x_*i*_ its context set of characters x/*M*. Specifically, for each training sample, we mask out a random 15% of the token positions for prediction. If the *i*-th token is chosen, it is replaced with (1) the *MASK* token 80% of the time (2) a random token 10% of the time (3) the unchanged i-th token 10% of the time. We use the output of the Transformer encoder, z to predict the likelihood of each amino acid a each missing token x_*i*_ using the following linear mapping:

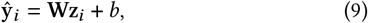

with learned matrix W ∈ ℝ^| *V* | ×*d*^, bias *b*, and DeepVHPPI output z_*i*_ ∈ ℝ^*d*×1^. *ŷ*_*i*_ is then fed through a softmax function to obtain class probabilities, *p* (*x*_*i*_ | *x*_/*M*_)

#### 3.3.2 SP: Structure Prediction Pretraining

Masked language model pretraining uses large amounts of unlabeled data to learn protein sequence semantics. However, a key aspect of whether two proteins interact or not is the structure of each protein. Obtaining structural information is slow and expensive, so we generally do not have structure information for all proteins from a novel virus. We leverage existing structural information to further pretrain DeepVHPPI and learn structure from sequence by predicting known structures. We consider three structure-based classification tasks: (1) Secondary Structure (SS) prediction (2) Contact prediction, and, (3) Homology prediction. Each task is explained below.

### Secondary Structure Prediction

Protein secondary structure is the three dimensional form of local segments of proteins. Each character in the sequence can be labeled by its secondary structure, which is one of | *C*| classes where *C* = {Helix, Strand, Other}. This results in a sequence tagging task where each input amino acid character x_*i*_ is mapped to a label *y*_*i*_ ∈ *C*. We predict the likelihood of each class for *x*_*i*_ using the following linear mapping:

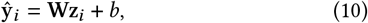

with learned matrix W ∈ℝ^| *C* | ×*d*^, bias *b*, and DeepVHPPI output z_*i*_ ∈ℝ^*d*×1^. *ŷ* is then fed through a softmax function to obtain class probabilities.

### Contact Prediction

Protein contact maps are a simplified depiction of the global 3D structure protein, where binary contact points indicated interactions in the 3D space. Contact prediction aims to predict the contact of each set of amino acid pairs in the sequence. Pair (*x*_*i*_, *x* _*j*_) of input amino acids from sequence **x** is mapped to a label *y*_*i j*_ ∈{0, 1} indicating whether or not the amino acids are physically close (< 8Å apart) to each other. To produce the contact likelihood of pair (*x*_*i*_, *x* _*j*_), we use the following formula which preserves non-directionality of contacts:

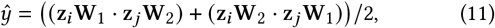

where {**W**_1_, **W**_2_} ∈ ℝ^*d*×*d*^. *ŷ* is then fed through a sigmoid function to obtain the contact probability.

### Remote Homology Detection

Remote homologues are pairs of proteins that share the same functional class, but have drastically different sequences. The goal of remote homology detection is predict the structural and functional class of a protein. Since proteins evolve, many proteins are structurally (and thus, functionally) similar, although their sequences are slightly different. Accurately predicting the homology of a protein would allow the model to group similar structural proteins together. We consider predicting the remote homology of a protein in terms of structural “fold” classes (e.g. Beta Barrel). This is a protein classification task where each input sequence x is mapped to a label *y* ∈*C*, where | *C*| = 1195 different possible protein folds. We use the designated *CLS* token from the DeepVHPPI to predict one of the | *C* | labels for a given sequence. We use a single linear layer mapping z_*i*_ to a | *C* | -dimensional vector:

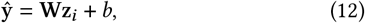

where **W** ∈ ℝ^| *C* | ×*d*^ is a projection matrix and z_*i*_ ∈ ℝ^*d*×1^ is the *CLS* token output vector from the Transformer. *ŷ* is fed through a softmax function to obtain class probabilities.

### Multi-task Training

For secondary structure and contact prediction, we use a binary cross-entropy loss, and for remote homoloy, we use cross entropy. We train all three structure prediction tasks simultaneously by sampling a new task uniformly in each gradient descent batch.

#### 3.3.3 Finetuning on V-H PPI Task

After MLM and SP pretraining, the final task is to finetune the model using the classifier in Section 3.2 on the known (i.e. experimentally validated) PPI data for previously known virus proteins. Once this model is trained, we can use it on pairs of Virus–Human interaction sequences where the virus was not used during training. The pretraining method on generic tasks learn structures of proteins which generalize well to unseen proteins.

## 4 CONNECTING TO RELATED WORK

### Protein-Protein Interaction Prediction

Many previous PPI works focus on developing intra-species interactions [10, 24, 30, 43, 56, 61, 65]. In other words, they would have one model for only Human– Human interactions and another model for only Yeast–Yeast interactions. Cross-species interaction prediction instead relates to where each protein in the interaction comes from a different species. Many works predict cross-species PPIs where the testing set contains proteins that are in the training set [6, 15, 40, 49, 62]. These methods do not reflect the real-world setting for a novel virus since we don’t have training proteins available for the virus. Additionally, PPI prediction methods generally perform much better for test pairs that share components with a training set than for those that do not [42]. Few works have focused on the more difficult task of cross-species interaction prediction where one of the protein species is completely unseen during training, which is what our work tackles. DeNovo [18] used an SVM for cross species interaction prediction. Yang et al. [68] introduce a deep learning embedding method combined with a random forest. Zhou et al. [71] improved DeNovo’s SVM for novel Virus–Human interaction.

### Protein Sequence Classification

Machine learning methods have achieved considerable results predicting properties of proteins that have yet to be experimentally validated by experimental studies. [35, 48] introduce multitask deep learning models for sequence labeling tasks such as SS prediction. [38, 52, 57] focus on methods of language model pretraining for generalizable representations of sequences. In particular, [52, 57] and [9] showed that self-supervised pretraining can produce protein representations that generalize across protein domains.

### Transformers

Transformers [63] obtained state-of-the-art results on several NLP tasks [17]. One problem with the vanilla Transformer model on token level inputs is that locality is not preserved. [5] used varying convolutional filters on characters at the word level and took the mean of the output to get a single vector representation for each word. Since proteins have no inherent “words” we use the convolutional output for each character as its local word. Instead of using character level inputs, word or byte-pair encodings can be used in order to preserve the local structure of words in text [59].

### Transfer Learning

Our work relates to several others in natural language processing where pretraining is used to transfer knowledge from both unlabeled and related labeled datasets [3, 34, 45]. Transfer learning is closely tied with few-shot learning [33, 53], which typically aims to use representations from prior tasks to generalize. Transformers are particularly well-fitted for transfer learning as their parallelizable architecture allows for fast pretraining on large datasets [17, 50]. It has been shown that this large-scale pretraining generalizes well enough for accurate few-shot learning [11].

## 5 EXPERIMENTAL SETUP AND RESULTS

### 5.1 Model Details and Evaluation Metrics

#### DeepVHPPI Variations and Details

We evaluate three variants of our model: (1) DeepVHPPI: this is the base model which uses no pretraining, only training on the target PPI task. (2) DeepVHPPI+MLM: this variant uses the language model pretraining and finetuning on the PPI task. (3) DeepVHPPI+MLM+SP: this uses both language model pretraining and supervised structure/family prediction pretraining before finetuning on the PPI task. We test both single task and multi-task models on the 3 SP tasks. We use the multi-task trained model for all PPI tasks. We train all models using a 12-layer transformer of size *d*=712 with *H*=8 attention heads.

For all training and testing, we clip protein sequences to 1024 length. For language model pretraining, we use a batch size of 1024, a linear warmup, and max learning rate of 1e-3. For all other tasks, we use a batch size of 16 with max learning rate of 1e-5. The language model is trained for 60 epochs, and all others are trained for 100. All models are trained with an Adam optimizer [31] and 10% dropout. Our models are implemented in PyTorch and we run each model on 4 NVIDIA Titan X GPUs. Masked language model pretraining (MLM) takes approximately 3 days, structure prediction (SP) pretraining takes 1 day, and the PPI task takes 3 days. Testing on ∼ 0.5M PPI pairs takes about 1 hour.

#### Metrics

For the supervised pretraining tasks, we use the metrics reported by previous work [52]. For the PPI task, we are largely focused on ranking interaction predictions based on probability, so we report two non-thresholding metrics: area under the ROC curve (AUROC), and area under the precision-recall curve (AUPR). We additionally report F1 scores where we consider thresholds [0.1,0.2,0.3,0.4,0.5,0.6,0.7,0.8,0.9]. As done in previous works, the results are selected on the best performing test epoch for each metric. For the SARS-CoV-2 dataset, we evaluate each metric for each virus protein individually since we are interested in the accuracy of predicting human interactions for specific virus proteins. The reported results are the mean value across all 25 virus proteins. For this dataset, we also report precision at 100 (P@100).

### 5.2 Pretraining Tasks (MLM and SP)

#### Datasets

For the masked language model (MLM) task, we train DeepVHPPI on all Swiss-Prot protein sequences [14]. Swiss-Prot is a collection of 562,253 manually reviewed, non-redundant protein sequences from 9,594 organisms. It includes most human and known virus proteins, allowing our model to learn the distribution of both types. We then further pretrain the model with structure pretraining (SP) tasks. For secondary structure prediction, the original data is from [32] and we report accuracy. For contact prediction, data is from [1]. We report precision of the *L*/5 most likely contacts (where *L* is the sequence length) for medium and long-range contacts. For remote homology prediction, the data is from [29], and we report accuracy. In all 3 tasks, we use the train/validation/test splits from [52].

#### Baselines

We compare our model against three deep learning methods: a vanilla Transformer, an LSTM [28], and ResNet [25], all run by [52]. Our DeepVHPPI uses the same Transformer model size (number of trainable parameters) as the vanilla transformer, and similar model size to the LSTM and ResNet. We do not compare to the pretrained models from [52] since we use a different pretraining dataset. We also compare our method against two baseline methods from [52]. “One-hot” uses one-hot feature inputs that are fed to simple classifiers such as an MLP or 2-layer ConvNet. “Alignment” uses sequence alignment features (BLAST or HHBlits), which are matrices that encode evolutionary information about the protein [55, 58].

#### Results

Here we investigate the benefits of the DeepVHPPI on the structure prediction tasks. Table 2 shows results on the three SP datasets. Our proposed DeepVHPPI performs as well or better than baseline methods, aside from alignment methods on SS prediction. Self supervised language modeling adds improvement over the base DeepVHPPI. Multi-task training with language model pretraining outperforms all other non-alignment methods. We do not evaluate the performance of the MLM task itself since better MLM performance does not always translate to better downstream task performance [57].

### 5.3 SARS-CoV-2–Human PPI Task

#### Dataset

While there may be no known Virus–Human interactions for a novel virus, there are many experimentally validated interactions for previous viruses. Our proposed approach is to train an interaction model on known V–H interactions (as indicated by solid lines in Fig 1), and then test on all possible V–H interactions for the novel virus proteins (as indicated by dotted lines in Fig 1). We explain our full training and testing setup below. A summary of the datasets used is provided in Table 1.

**Table 1:** Datasets: For each category of training: Language Model (LM), Intermediate (SP) and PPI, we provide the dataset output type and training/validation/test set sizes. *L* represents the sequence length, and | *V*| represents the vocabulary size.

| Dataset | Category | Output Shape | Total | Train | Valid | Test |
| --- | --- | --- | --- | --- | --- | --- |
| Swiss-Prot | MLM | $L \times \vert V\vert $ | 562,280 | 562,280 | N/A | N/A |
| Secondary Structure | SP | $L \times 3$ | 11,361 | 8,678 | 2,170 | 513 |
| Contact | SP | $L \times L$ | 25,563 | 25,299 | 224 | 40 |
| Homology | SP | $1195 \times 1$ | 13,766 | 12,312 | 736 | 718 |
| SARS-CoV-2 | PPI | $1 \times 1$ | 815,279 | 199,346 | 49,836 | 610,950 |
| H1N1 [71] | PPI | $1 \times 1$ | 22,291 | 21,910 | N/A | 381 |
| Ebola [71] | PPI | $1 \times 1$ | 22,982 | 22,682 | N/A | 300 |

**Table 2:** Structure prediction (SP) pretraining task results. For SS and Homology, accuracy is reported. For Contact, precision at *L* 5 for for medium and long-range contacts is reported.

| Method | SS | Contact | Homology |
| --- | --- | --- | --- |
| One-hot [52] | 0.69 | 0.29 | 0.09 |
| Alignment [52] | **0.80** | 0.64 | 0.09 |
| ResNet [52] | 0.70 | 0.20 | 0.10 |
| LSTM [52] | 0.71 | 0.19 | 0.12 |
| Transformer [52] | 0.70 | 0.32 | 0.09 |
| DeepVHPPI | 0.70 | 0.51 | 0.12 |
| DeepVHPPI + MLM | 0.71 | 0.58 | 0.22 |
| DeepVHPPI (multi-task) | 0.64 | 0.53 | 0.13 |
| DeepVHPPI + MLM (multi-task) | 0.71 | **0.70** | **0.38** |

##### Training Data

We use the V–H dataset from Yang et al. [66], which is based on data from the Host-Pathogen interaction Database (HPIDB; version 3.0) [2]. Note this does only contains Host-Pathogen interactions, and not contain Human-Human interactions. This dataset processed interactions from large-scale mass spectrometry experiments, resulting in 22,653 experimentally verified host-virus PPIs as a positive sample set. The authors chose negative pairs based on the ‘Dissimilarity-Based Negative Sampling’ [18]. The selected ratio of positive to negative samples is 1:10. Following Yang et al., we use the same training set (80%) and an independent validation set (20%) for model training and hyperparameter selection, respectively.

##### Testing Data

For the novel virus testing dataset, we use the 13,947 known SARS-CoV-2–Human interactions from the BioGRID database (Coronavirus version 4.1.190) [41]. All BioGRID interactions are experimentally validated, many from [21]. Considering all 20,365 SwissProt human proteins, we label all other pairs from the total space of 20,365* 26 to be non-interacting (a total of 529,490). It is important to note that the labeled “negative” samples contain many pairs of proteins that interact but are not known to do so. This can result in an overestimation of the false positive rate [20].

#### Baselines

We compare our method to the pretrained Embedding+RF method from Yang et al. [68], which uses Doc2Vec to embed protein sequences and then a random forest to classify pairs. To the best of our knowledge, no other methods provide code or a browser service to run novel protein interactions.

#### Results

Table 3 shows our results for SARS-CoV-2. Across most metrics, our method DeepVHPPI outperforms the baseline method. MLM and SP pretraining help generalize to our target PPI task, where the non-pretrained models aren’t as accurate. We note that the testing dataset is highly imbalanced. In other words, most interactions (99.7%) are negative, or non-interacting. Thus, the AUROC metric is not indicative of good results. We turn our attention to the AUPR metrics, where our method performs the best. This confirms our hypothesis that pretraining on large protein datasets learn evolutionary structures of proteins which generalize well to unseen proteins. This is a promising result not only for SARS-CoV-2, but for potential future novel viruses.

**Table 3:** Human and SARS-CoV-2 Interaction Predictions. Each metric is reported as the mean across all virus proteins. Best results are reported in bold.

| Method | AUROC | AUPR | F1(%) | P@100 |
| --- | --- | --- | --- | --- |
| Embedding+RF [68] | 0.748 | 0.071 | **0.116** | 0.126 |
| DeepVHPPI | 0.740 | 0.065 | 0.104 | 0.129 |
| DeepVHPPI+MLM | 0.751 | 0.070 | 0.110 | 0.147 |
| DeepVHPPI+MLM+SP | **0.753** | **0.076** | 0.114 | **0.151** |

### 5.4 Other Virus–Host PPI Tasks (H1N1 and Ebola)

#### Datasets

In our SARS-CoV-2 dataset, we explain a testing scenario where we have no knowledge of the true V–H interactions, resulting in a large possible interaction space (all possible | *P*_*ν*_| × | *P*_*h*_| interactions). Zhou et al. [71] created V–H datasets where they hand selected the negative interactions based on the known positives, making sure that they had an even positive/negative split. While this setup is unrealistic for a true novel virus (because we don’t know which ones are positive), we compare to their results to show the strength of our method. Zhou et a. provide two H–V datasets, H1N1 and Ebola.

In the H1N1 dataset, the training set contains 10,955 true PPIs between human and any virus except H1N1 virus, plus an equal amount (10,955) negative interaction samples. The testing set contains 381 true PPIs between human and H1N1 virus, and 381 negative interactions. Similarly, in the Ebola dataset, the training set contains 11,341 true PPIs between human and any virus except Ebola virus, plus an equal amount (11,341) negative interaction samples. The testing set contains 150 true PPIs between human and Ebola virus, and 150 negative interactions.

#### Baseline

We compare to the SVM baseline from Zhou et al. [71], which showed that their method was the state-of-the-art.

#### Results

Table 4, shows the results from the Zhou et al. datasets. For both the H1N1 and Ebola datasets, we can see that our method outperforms the previous state-of-the-art. While we believe this dataset is not indicative of a real novel virus setting since the test set negatives are hand-selected, we can use it to compare different methods. Since there is an even positive/negative testing split, AUROC is a good metric to compare methods, and we can see that across both novel viruses, our method outperforms the SVM. We see notable performance increase using LMT and SP pretraining.

**Table 4:** Virus–Human PPI Tasks from Zhou et al. [71]. Best results are in bold. “-” indicates the metric was not reported.

| Method | H1N1 |  |  | Ebola |  |  |
| --- | --- | --- | --- | --- | --- | --- |
|  | AUROC | AUPR | F1(%) | AUROC | AUPR | F1(%) |
| SVM [71] | 0.886 | - | 76.2 | 0.867 | - | 76.0 |
| DeepVHPPI | 0.903 | 0.898 | 84.4 | 0.912 | 0.953 | 86.0 |
| DeepVHPPI+MLM | 0.908 | 0.903 | 85.5 | 0.953 | 0.961 | **90.4** |
| DeepVHPPI+MLM+SP | **0.945** | **0.948** | **86.5** | **0.968** | **0.974** | 89.6 |

##### 5.5 Additional PPI Experiments

We also report the results for our model on two extra baseline datasets. The first is the Virus–Host datasets from Barman et al. [6]. Although the testing dataset is not for a completely unseen test virus, we see our method outperforms the baselines. In Table 5, we see that our method outperforms previous methods including an SVM and random forest.

**Table 5:** Virus–Human PPI Tasks from Barman et al. [6]. Best results are in bold. “-” indicates the metric was not reported.

| Method | AUROC | AUPR | F1 |
| --- | --- | --- | --- |
| SVM [71] | 0.858 | - | 79.2 |
| Embedding + RF [68] | 0.871 | - | 79.8 |
| DeepVHPPI+UPT+SPT | **0.886** | **0.806** | **80.6** |

The second is the SLiMs dataset from Eid et al. [18]. This dataset was constructed specifically to evaluate how well the model learns Short Linear Motifs (SLiMs) that are transferable across train/test splits. In Table 6, we see that our model is significantly better than the baselines. We hypothesize that this improvement is in part due to the convolutional motif finder layers of DeepVHPPI.

**Table 6:** SLiM PPI Tasks from Eid et al. [18]. Best results are in bold. “-” indicates the metric was not reported.

| Method | AUROC | AUPR | F1 |
| --- | --- | --- | --- |
| DeNovo [18] | - | - | 81.9 |
| SVM [71] | 0.897 | - | 84.2 |
| DeepVHPPI+UPT+SPT | **0.989** | **0.991** | **95.9** |

##### 5.6 Sensitivity Analysis using Known H-V Interactions

There exist two important situations we need to predict V-H PPI for virus protein with new sequences. (I) The first case is when a new virus is discovered, the protein interactions are unknown. (II) The second case is when the virus is known, and some interactions have been validated, but many are missing. Our above experiments have mostly focused on the first case. Here we experiment and analyze when some of the virus proteins are known. This setup can be used to expand the initial set of experimental interactions, resulting in a more complete interactome.

On the SARS-CoV-2–Human PPI task, we simulate a new setup to evaluate our framework across these two settings. We generate five simulated settings where we vary the percentage of total SARS-CoV-2–Human interactions to be used as training data. We then test on the remaining SARS-CoV-2–Human interactions. The training percentages we use are 0%,20%,40%,60%,80%. We also consider training on all known Human–Human (H–H) interactions from the BioGrid database. Note that we are not using any other virus proteins in these simulations. Fig. 4 shows the results across each of the five settings for four DeepVHPPI model variations and a deep learning baseline from Rao et al. [52], which uses MLM pretraining.

**Figure 4:**
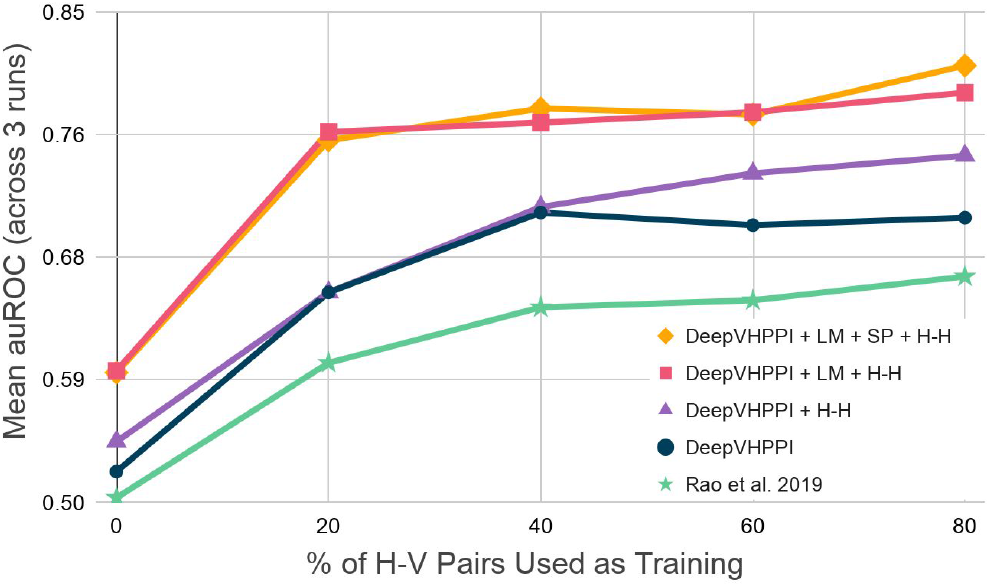
Sensitivity analysis on Human–SARS-CoV-2 V-H-PPI Predictions. The X-axis shows simulated training settings, where we assume there exist some (varying) portion of SARS-CoV-2 proteins in the training data. We can see that pretraining methods (LM and SPT) give substantial increases over cases without the pretraining methods. This indicates that transfer learning can help on novel virus protein interaction prediction.

First, across all training settings, we can see that DeepVHPPI models outperform the baseline [52]. Across all settings, adding the transfer learning pretraining tasks helps significantly. Most importantly, we can see that in the case I (0%) and case II (20%) settings, MLM and SP pretraining help generalize to the target task, where the non-pretrained models fail to be able to classify well. Additionally, adding H–H interactions helps generalize for predicting novel virus sequence interactions.

##### 5.7 Mutation Validation Analysis on SARS-CoV-2 Spike

Understanding how a classifier changes its predictions based on small perturbations in the sequence is an important tool for interpreting deep learning models [39]. A commonly used strategy is to replace elements of a sequence with mutations, and look at how the classifier degrades. This method is cheap to evaluate but comes at a significant drawback. Samples where a subset of the features are replaced come from a different distribution. Therefore, this approach clearly violates one of the key assumptions in machine learning: the training and evaluation data come from the same distribution. Without re-training, it is unclear whether the degradation in prediction performance comes from the distribution shift or because the features that were removed are truly informative. For this reason we decide to verify how much information can be removed before accuracy of a retrained model breaks down completely.

We used the deep mutational scan data from Starr et al. [60] to analyze the effectiveness of DeepVHPPI the accurately model binding affinity of virus mutations. This dataset contains 105,526 mutated SARS-CoV-2 Spike receptor binding domain sequences, with the corresponding Human ACE2 dissociation constant. The Spearman correlation between DeepVHPPI binding prediction and the in-vitro dissociation constant was 0.110 (pvalue=5.3e-235). DeepVHPPI was trained without any knowledge of the Spike protein, only previous virus and host interactions from HPIDB.

Furthermore, our method allows for an easy way to rapidly test new mutations. Rather than experimentally checking all possible mutations to see which ones reduce interaction binding, we can computationally introduce mutations and observe how the predicted output changes. We show an example of this for a receptor-binding domain subsequence of the SARS-CoV-2 Spike protein when binding to the human ACE2 protein in Fig. 5. We can observe specific locations, such as the first “K” amino acid in the virus sequence where mutating the amino acid will reduce the interaction prediction. There are also other locations where mutations can *increase* interaction binding, which may explain how certain viruses are able to mutate and infect humans more easily.

**Figure 5:**
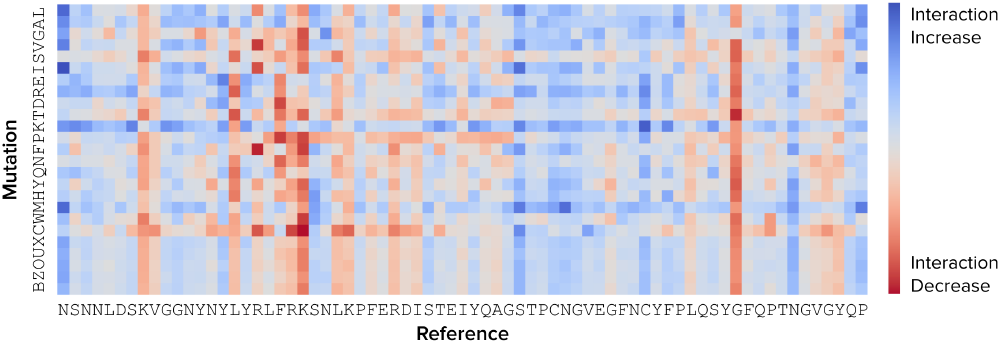
Mutation map on SARS-CoV-2 Spike when interacting with Human ACE2. Here we show how the predicted interaction score changes when inducing the mutation in the Y-axis for the original amino acid in the X-axis. This example is from the receptor-binding domain in the SARS-CoV-2 Spike protein, and the output shows the predicted difference in interaction score from the original reference amino acid. For example, changing the second reference “K” results in an interaction decrease.

## 6 CONCLUSION

Computational methods predicting protein-protein interactions (PPIs) can play an important role in understanding a novel virus that threatens widespread public health. Most previous methods are developed for intra-species interactions, and do not generalize to novel viruses. In this paper, we introduce DeepVHPPI, a novel deep learning architecture that uses a transfer learning approach for PPI prediction between a novel virus and a host. We show that our method can help accurately predict Virus–Host interactions early on in the virus discovery and experimentation pipeline. This can help biologists better understand how the virus attacks the human body, allowing researchers to potentially develop effective drugs more quickly. By providing a computational model for interaction prediction, we hope this will accelerate experimental efforts to define a reliable network of Virus–Host protein interactions. While this work is focused on SARS-CoV-2, H1N1, and Ebola, our framework is applicable for any virus. In the case of a future novel virus, our framework will be able to rapidly predict protein-protein interaction predictions.

## Acknowledgements

This work was partly supported by the National Science Foundation under NSF CAREER award No. 1453580 to Y.Q, as well as the Google Cloud COVID-19 Credits Program. Any opinions, findings and conclusions or recommendations expressed in this material are those of the author(s) and do not necessarily reflect those of the National Science Foundation.

## Footnotes

1 Mechanisms by which virus infection leads to disease in the target host

## REFERENCES

[1] Mohammed AlQuraishi. End-to-end differentiable learning of protein structure. Cell systems, 8(4):292–301, 2019.

[2] Mais G Ammari, Cathy R Gresham, Fiona M McCarthy, and Bindu Nanduri. Hpidb 2.0: a curated database for host–pathogen interactions. Database, 2016, 2016.

[3] Rie Kubota Ando and Tong Zhang. A framework for learning predictive structures from multiple tasks and unlabeled data. Journal of Machine Learning Research, 6(Nov):1817–1853, 2005.

[4] Jimmy Lei Ba, Jamie Ryan Kiros, and Geoffrey E Hinton. Layer normalization. arXiv preprint 1607.06450, 2016.

[5] Alexei Baevski, Sergey Edunov, Yinhan Liu, Luke Zettlemoyer, and Michael Auli. Cloze-driven pretraining of self-attention networks. arXiv preprint 1903.07785, 2019.

[6] Ranjan Kumar Barman, Sudipto Saha, and Santasabuj Das. Prediction of interactions between viral and host proteins using supervised machine learning methods. PloS one, 9(11):e112034, 2014.

[7] Asa Ben-Hur and William Stafford Noble. Kernel methods for predicting protein– protein interactions. Bioinformatics, 21(Suppl_1):i38–i46, 2005.

[8] Yoshua Bengio, Réjean Ducharme, Pascal Vincent, and Christian Jauvin. A neural probabilistic language model. Journal of machine learning research, 3(Feb):1137–1155, 2003.

[9] Tristan Bepler and Bonnie Berger. Learning protein sequence embeddings using information from structure. arXiv preprint 1902.08661, 2019.

[10] Anne-Florence Bitbol. Inferring interaction partners from protein sequences using mutual information. PLoS computational biology, 14(11):e1006401, 2018.

[11] Tom B. Brown, Benjamin Mann, Nick Ryder, Melanie Subbiah, Jared Kaplan, Prafulla Dhariwal, Arvind Neelakantan, Pranav Shyam, Girish Sastry, Amanda Askell, Sandhini Agarwal, Ariel Herbert-Voss, Gretchen Krueger, Tom Henighan, Rewon Child, Aditya Ramesh, Daniel M. Ziegler, Jeffrey Wu, Clemens Winter, Christopher Hesse, Mark Chen, Eric Sigler, Mateusz Litwin, Scott Gray, Benjamin Chess, Jack Clark, Christopher Berner, Sam McCandlish, Alec Radford, Ilya Sutskever, and Dario Amodei. Language models are few-shot learners, 2020.

[12] Ronan Collobert, Jason Weston, Léon Bottou, Michael Karlen, Koray Kavukcuoglu, and Pavel Kuksa. Natural language processing (almost) from scratch. Journal of machine learning research, 12(Aug):2493–2537, 2011.

[13] Qian Cong, Ivan Anishchenko, Sergey Ovchinnikov, and David Baker. Protein interaction networks revealed by proteome coevolution. Science, 365(6449):185–189, 2019.

[14] UniProt Consortium. Uniprot: a worldwide hub of protein knowledge. Nucleic acids research, 47(D1):D506–D515, 2019.

[15] Guangyu Cui, Chao Fang, and Kyungsook Han. Prediction of protein-protein interactions between viruses and human by an svm model. In BMC bioinformatics, volume 13, p. S5. Springer, 2012.

[16] Norman E Davey, Gilles Travé, and Toby J Gibson. How viruses hijack cell regulation. Trends in biochemical sciences, 36(3):159–169, 2011.

[17] Jacob Devlin, Ming-Wei Chang, Kenton Lee, and Kristina Toutanova. Bert: Pretraining of deep bidirectional transformers for language understanding. arXiv preprint 1810.04805, 2018.

[18] Fatma-Elzahraa Eid, Mahmoud ElHefnawi, and Lenwood S Heath. Denovo: virus-host sequence-based protein–protein interaction prediction. Bioinformatics, 32(8):1144–1150, 2016.

[19] Stanley Fields and Ok-kyu Song. A novel genetic system to detect protein–protein interactions. Nature, 340(6230):245–246, 1989.

[20] Shawn M Gomez, William Stafford Noble, and Andrey Rzhetsky. Learning to predict protein–protein interactions from protein sequences. Bioinformatics, 19(15):1875–1881, 2003.

[21] David E Gordon, Gwendolyn M Jang, Mehdi Bouhaddou, Jiewei Xu, Kirsten Obernier, Kris M White, Matthew J O’Meara, Veronica V Rezelj, Jeffrey Z Guo, Danielle L Swaney, et al. A sars-cov-2 protein interaction map reveals targets for drug repurposing. Nature, pp. 1–13, 2020.

[22] Yanzhi Guo, Lezheng Yu, Zhining Wen, and Menglong Li. Using support vector machine combined with auto covariance to predict protein–protein interactions from protein sequences. Nucleic acids research, 36(9):3025–3030, 2008.

[23] Tobias Hamp and Burkhard Rost. Evolutionary profiles improve protein–protein interaction prediction from sequence. Bioinformatics, 31(12):1945–1950, 2015.

[24] Somaye Hashemifar, Behnam Neyshabur, Aly A Khan, and Jinbo Xu. Predicting protein–protein interactions through sequence-based deep learning. Bioinformatics, 34(17):i802–i810, 2018.

[25] Kaiming He, Xiangyu Zhang, Shaoqing Ren, and Jian Sun. Deep residual learning for image recognition. In Proceedings of the IEEE conference on computer vision and pattern recognition, pp. 770–778, 2016.

[26] Dan Hendrycks and Kevin Gimpel. Gaussian error linear units (gelus). arXiv preprint 1606.08415, 2016.

[27] Yuen Ho, Albrecht Gruhler, Adrian Heilbut, Gary D Bader, Lynda Moore, Sally-Lin Adams, Anna Millar, Paul Taylor, Keiryn Bennett, Kelly Boutilier, et al. Systematic identification of protein complexes in saccharomyces cerevisiae by mass spectrometry. Nature, 415(6868):180–183, 2002.

[28] Sepp Hochreiter and Jürgen Schmidhuber. Long short-term memory. Neural computation, 9(8):1735–1780, 1997.

[29] Jie Hou, Badri Adhikari, and Jianlin Cheng. Deepsf: deep convolutional neural network for mapping protein sequences to folds. Bioinformatics, 34(8):1295–1303, 2018.

[30] Kalyani B Karunakaran, N Balakrishnan, and Madhavi K Ganapathiraju. Interactome of sars-cov-2/ncov19 modulated host proteins with computationally predicted ppis, 2020.

[31] Diederik P Kingma and Jimmy Ba. Adam: A method for stochastic optimization. arXiv preprint 1412.6980, 2014.

[32] Michael Schantz Klausen, Martin Closter Jespersen, Henrik Nielsen, Kamilla Kjaergaard Jensen, Vanessa Isabell Jurtz, Casper Kaae Soenderby, Morten Otto Alexander Sommer, Ole Winther, Morten Nielsen, Bent Petersen, et al. Netsurfp-2.0: Improved prediction of protein structural features by integrated deep learning. Proteins: Structure, Function, and Bioinformatics, 87(6):520–527, 2019.

[33] Zhenguo Li, Fengwei Zhou, Fei Chen, and Hang Li. Meta-sgd: Learning to learn quickly for few-shot learning. arXiv preprint 1707.09835, 2017.

[34] Dekang Lin and Xiaoyun Wu. Phrase clustering for discriminative learning. In Proceedings of the Joint Conference of the 47th Annual Meeting of the ACL and the 4th International Joint Conference on Natural Language Processing of the AFNLP: Volume 2-Volume 2, pp. 1030–1038. Association for Computational Linguistics, 2009.

[35] Zeming Lin, Jack Lanchantin, and Yanjun Qi. Must-cnn: a multilayer shift-and-stitch deep convolutional architecture for sequence-based protein structure prediction. In Thirtieth AAAI conference on artificial intelligence, 2016.

[36] Shawn Martin, Diana Roe, and Jean-Loup Faulon. Predicting protein–protein interactions using signature products. Bioinformatics, 21(2):218–226, 2005.

[37] Tomas Mikolov, Ilya Sutskever, Kai Chen, Greg S Corrado, and Jeff Dean. Distributed representations of words and phrases and their compositionality. In Advances in neural information processing systems, pp. 3111–3119, 2013.

[38] Seonwoo Min, Seunghyun Park, Siwon Kim, Hyun-Soo Choi, and Sungroh Yoon. Pre-training of deep bidirectional protein sequence representations with structural information, 2019.

[39] John X Morris, Eli Lifland, Jack Lanchantin, Yangfeng Ji, and Yanjun Qi. Reevaluating adversarial examples in natural language. arXiv preprint 2004.14174, 2020.

[40] Esmaeil Nourani, Farshad Khunjush, and Saliha Durmuş. Computational approaches for prediction of pathogen-host protein-protein interactions. Frontiers in microbiology, 6:94, 2015.

[41] Rose Oughtred, Chris Stark, Bobby-Joe Breitkreutz, Jennifer Rust, Lorrie Boucher, Christie Chang, Nadine Kolas, Lara O’Donnell, Genie Leung, Rochelle McAdam, et al. The biogrid interaction database: 2019 update. Nucleic acids research, 47(D1):D529–D541, 2019.

[42] Yungki Park and Edward M Marcotte. Flaws in evaluation schemes for pair-input computational predictions. Nature methods, 9(12):1134, 2012.

[43] Florencio Pazos and Alfonso Valencia. Similarity of phylogenetic trees as indicator of protein–protein interaction. Protein engineering, 14(9):609–614, 2001.

[44] Jeffrey Pennington, Richard Socher, and Christopher D Manning. Glove: Global vectors for word representation. In Proceedings of the 2014 conference on empirical methods in natural language processing (EMNLP), pp. 1532–1543, 2014.

[45] Matthew E Peters, Waleed Ammar, Chandra Bhagavatula, and Russell Power. Semi-supervised sequence tagging with bidirectional language models. arXiv preprint 1705.00108, 2017.

[46] EM Phizicky and S. Fields. Protein-protein interactions: methods for detection and analysis. Microbiol Rev., 59(1):94–123, 1995.

[47] Sylvain Pitre, Mohsen Hooshyar, Andrew Schoenrock, Bahram Samanfar, Matthew Jessulat, James R Green, Frank Dehne, and Ashkan Golshani. Short co-occurring polypeptide regions can predict global protein interaction maps. Scientific reports, 2:239, 2012.

[48] Yanjun Qi, Merja Oja, Jason Weston, and William Stafford Noble. A unified multitask architecture for predicting local protein properties. PloS one, 7(3):e32235, 2012.

[49] Yanjun Qi, Oznur Tastan, Jaime G Carbonell, Judith Klein-Seetharaman, and Jason Weston. Semi-supervised multi-task learning for predicting interactions between hiv-1 and human proteins. Bioinformatics, 26(18):i645–i652, 2010.

[50] Alec Radford, Jeffrey Wu, Rewon Child, David Luan, Dario Amodei, and Ilya Sutskever. Language models are unsupervised multitask learners, 2019.

[51] Prajit Ramachandran, Niki Parmar, Ashish Vaswani, Irwan Bello, Anselm Levskaya, and Jonathon Shlens. Stand-alone self-attention in vision models. arXiv preprint 1906.05909, 2019.

[52] Roshan Rao, Nicholas Bhattacharya, Neil Thomas, Yan Duan, Xi Chen, John Canny, Pieter Abbeel, and Yun S Song. Evaluating protein transfer learning with tape. arXiv preprint 1906.08230, 2019.

[53] Sachin Ravi and Hugo Larochelle. Optimization as a model for few-shot learning, 2016.

[54] Emma Redhead and Timothy L Bailey. Discriminative motif discovery in dna and protein sequences using the deme algorithm. BMC bioinformatics, 8(1):385, 2007.

[55] Michael Remmert, Andreas Biegert, Andreas Hauser, and Johannes Söding. Hhblits: lightning-fast iterative protein sequence searching by hmm-hmm alignment. Nature methods, 9(2):173, 2012.

[56] Florian Richoux, Charlène Servantie, Cynthia Borès, and Stéphane Téletchéa. Comparing two deep learning sequence-based models for protein-protein interaction prediction. arXiv preprint 1901.06268, 2019.

[57] Alexander Rives, Siddharth Goyal, Joshua Meier, Demi Guo, Myle Ott, C Lawrence Zitnick, Jerry Ma, and Rob Fergus. Biological structure and function emerge from scaling unsupervised learning to 250 million protein sequences. bioRxiv, p. 622803, 2019.

[58] Alejandro A Schä ffer, L Aravind, Thomas L Madden, Sergei Shavirin, John L Spouge, Yuri I Wolf, Eugene V Koonin, and Stephen F Altschul. Improving the accuracy of psi-blast protein database searches with composition-based statistics and other refinements. Nucleic acids research, 29(14):2994–3005, 2001.

[59] Rico Sennrich, Barry Haddow, and Alexandra Birch. Neural machine translation of rare words with subword units. arXiv preprint 1508.07909, 2015.

[60] Tyler N Starr, Allison J Greaney, Sarah K Hilton, Daniel Ellis, Katharine HD Crawford, Adam S Dingens, Mary Jane Navarro, John E Bowen, M Alejandra Tortorici, Alexandra C Walls, et al. Deep mutational scanning of sars-cov-2 receptor binding domain reveals constraints on folding and ace2 binding. Cell, 182(5):1295–1310, 2020.

[61] Tanlin Sun, Bo Zhou, Luhua Lai, and Jianfeng Pei. Sequence-based prediction of protein protein interaction using a deep-learning algorithm. BMC bioinformatics, 18(1):1–8, 2017.

[62] Oznur Tastan, Yanjun Qi, Jaime G Carbonell, and Judith Klein-Seetharaman. Prediction of interactions between hiv-1 and human proteins by information integration, 2009.

[63] Ashish Vaswani, Noam Shazeer, Niki Parmar, Jakob Uszkoreit, Llion Jones, Aidan N Gomez, Ł ukasz Kaiser, and Illia Polosukhin. Attention is all you need. In Advances in neural information processing systems, pp. 5998–6008, 2017.

[64] Christian von Mering, Roland Krause, Berend Snel, Michael Cornell, Stephen G. Oliver, Stanley Fields, and Peer Bork. Comparative assessment of large-scale data sets of protein-protein interactions. Nature, 417(6887):399–403, 2002.

[65] Lei Wang, Hai-Feng Wang, San-Rong Liu, Xin Yan, and Ke-Jian Song. Predicting protein-protein interactions from matrix-based protein sequence using convolution neural network and feature-selective rotation forest. Scientific reports, 9(1):1–12, 2019.

[66] Kevin K Yang, Zachary Wu, and Frances H Arnold. Machine-learning-guided directed evolution for protein engineering. Nature methods, 16(8):687–694, 2019.

[67] Lei Yang, Jun-Feng Xia, and Jie Gui. Prediction of protein-protein interactions from protein sequence using local descriptors. Protein and Peptide Letters, 17(9):1085–1090, 2010.

[68] Xiaodi Yang, Shiping Yang, Qinmengge Li, Stefan Wuchty, and Ziding Zhang. Prediction of human-virus protein-protein interactions through a sequence embedding-based machine learning method. Computational and structural biotechnology journal, 18:153–161, 2020.

[69] Zhu-Hong You, Lin Zhu, Chun-Hou Zheng, Hong-Jie Yu, Su-Ping Deng, and Zhen Ji. Prediction of protein-protein interactions from amino acid sequences using a novel multi-scale continuous and discontinuous feature set. In BMC bioinformatics, volume 15, p. S9. Springer, 2014.

[70] Shao-Wu Zhang and Ze-Gang Wei. Some remarks on prediction of proteinprotein interaction with machine learning. Medicinal Chemistry, 11(3):254–264, 2015.

[71] Xiang Zhou, Byungkyu Park, Daesik Choi, and Kyungsook Han. A generalized approach to predicting protein-protein interactions between virus and host. BMC genomics, 19(6):568, 2018.

